# The Rayleigh Quotient and Contrastive Principal Component Analysis I

**DOI:** 10.1101/2025.11.19.689125

**Authors:** Maria Carilli, Kayla Jackson, Lior Pachter

**Affiliations:** Division of Biology and Biological Engineering, California Institute of Technology, Pasadena, CA, USA; Department of Computing and Mathematical Sciences, California Institute of Technology, Pasadena, CA, USA

## Abstract

Contrastive learning methods can be powerful tools for genomics, enabling the identification of signals in an experiment via dimension reduction while reducing noise using a control. One such popular approach is contrastive PCA, which, despite being used in a variety of settings, does not scale to large datasets. We show that the contrastive PCA objective is an approximation of a Rayleigh quotient, analogous in form to Fisher’s linear discriminant analysis and the common spatial patterns method. The Rayleigh quotient is *ρ*PCA, satisfies numerous desirable properties, and provides an interpretable form of dimension reduction via generalized eigenvectors. We demonstrate that *ρ*PCA is more accurate than contrastive PCA and much more efficient. We also show how it can be used not only for dimension reduction of data with respect to a control, but also for contrasting conditions via an analysis of single-nucleus transcriptomics data. Finally, we discuss probabilistic interpretations of *ρ*PCA that provide further insight into its effective performance.

## Introduction

Principal Component Analysis (PCA) refers to the identification of a sequence of subspaces of a Euclidean vector space that provide optimal low-dimensional linear approximations to a high-dimensional dataset (Pearson, 1901; Hotelling, 1933). Formally, given centered data vectors *x*, …, *x* ∈ ℝ^*d*^ with sample covariance matrix 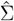, PCA seeks unit vectors *v*_1_, …, *v*_*k*_ that maximize

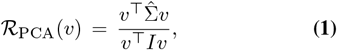

The vectors 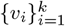 that maximize *ℛ*_PCA_(*v*) are the eigen-vectors of 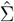, and the corresponding eigenvalues represent the variances explained along each direction. The *k*-dimensional subspace spanned by the first *k* eigenvectors maximizes the total projected variance and equivalently minimizes the mean-squared reconstruction error among all *k*-dimensional linear subspaces.

The quantity *ℛ*_PCA_(*v*) is a special case of the Rayleigh quotient, which, for two square matrices *A* and *B*, is defined as

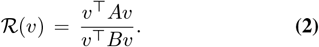

When both *A* and *B* are symmetric and *B* is positive definite, the maxima of (2) satisfy the generalized eigenvalue problem *Av* = *λBv* (see Methods; Horn and Johnson (2012)).PCA (1) corresponds to the special case where 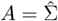 and *B* = *I*. The maximization of (2) via a generalized eigenvalue problem leads to numerous desirable properties for Rayleigh quotient solutions (see Methods), many of which are crucial in applications and contribute to PCA being a popular dimension-reduction method.

Many other dimension-reduction and signal-extraction problems can be formulated as a Rayleigh quotient maximization. For example, Fisher linear discriminant analysis (LDA; Fisher (1936); Mika et al. (1999)) maximizes the Rayleigh quotient

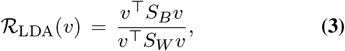

where *S*_*B*_ is a between-class scatter matrix and *S*_*W*_ is a within-class scatter matrix. Maximizing *ℛ*_LDA_(*v*) yields directions that best separate classes by maximizing inter-class variance while minimizing intra-class variance.

More recently, contrastive PCA (cPCA; Abid et al. (2018)) has been proposed as a tool to identify patterns present in one dataset but not another. The method seeks components that exhibit high variance in a target dataset while exhibiting low variance in a background dataset. The original publication defines cPCA as the eigendecomposition of a contrastive “covariance” matrix,

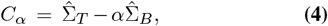

where 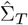 and 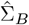 are the sample target and background co-variance matrices, respectively, and *α >* 0 is a tunable contrast parameter. Note that while the sum of covariance matrices is always a covariance matrix, the difference of co-variance matrices may not be positive semidefinite; *C*_*α*_ can have negative eigenvalues. Multiple subsequent publications have extended the original cPCA method, e.g. (Boileau et al., 2020; Li et al., 2024).

We observe that the cPCA objective (4) can be viewed as an approximate heuristic for solving the Rayleigh quotient problem

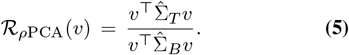

In particular, the cPCA objective (4) can be derived as a Taylor approximation (under a strong assumption) of the *ρ*PCA objective (5) (see Methods). Moreover, the *ρ*PCA objective (5) effectively identifies directions for which variance is maximized in the target data while it is minimized in the background data (Fig. 1). Unlike the cPCA formulation, maximizing *ℛ*_*ρ*PCA_(*v*) directly yields the exact solution to (5) without requiring tuning of the parameter *α* (see Methods). Furthermore, the resulting components are interpretable, computationally efficient to compute, and generalize naturally to settings with multiple target or background covariances.

**Fig. 1.**
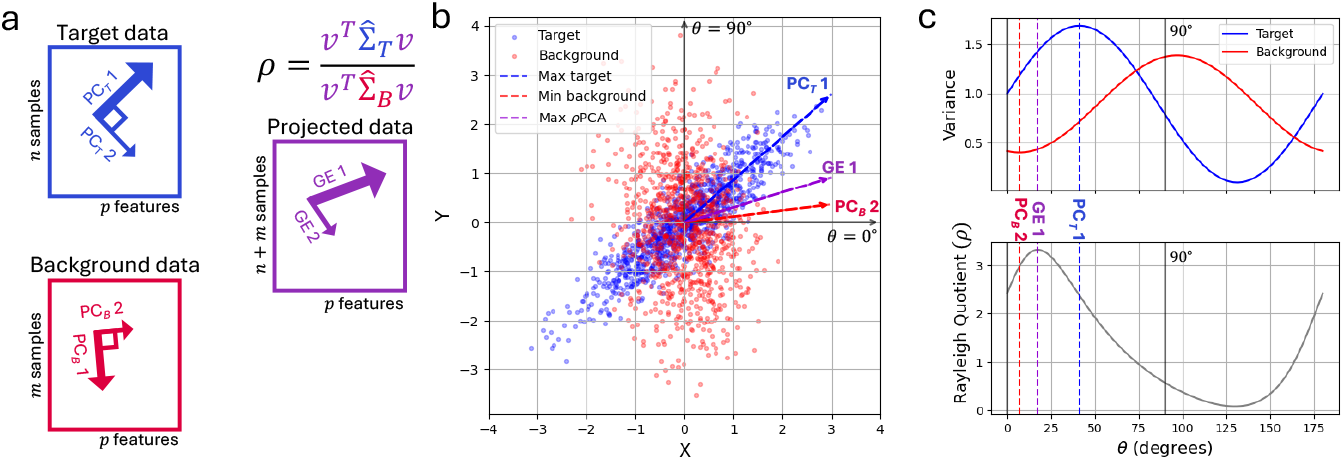
*ρ*PCA finds directions that maximize target variance while minimizing background variance. **a**, *ρ*PCA is useful in a setting in which there are target datasets of interest with axes of interesting variation and background datasets with different axes of variation that should be removed before analysis. In the following example, the sample size *n = m = 1,000* and the feature size *p=2*. **b**, *ρ*PCA (black dashed line) finds the direction that maximizes the ratio of the target variance (direction of maximum target variance, dashed blue line) while minimizing the background variance (direction of minimum background variance, red dashed line). **c**, Variance of target (blue) and background (red) data projected on lines that pass through the origin from 0°to 180°, with the ratio of target to background variance (Rayleigh Quotient) shown below. Dashed lines correspond to the directions from **b**.

## Results

Given the heuristics and approximations underlying cPCA, we hypothesized that *ρ*PCA would be more effective at finding a projection that maximizes variance in a target dataset while minimizing variance in a background dataset. In a setting where the target dataset and background dataset both share variance in two principal directions, albeit differ in a third direction where target variance is much higher than the background (Fig. 2a), *ρ*PCA can find an effective projection displaying the variation unique to the target data in GE 1 (Fig. 2b), while cPCA cannot (Fig. 2c). Notably, the failure of cPCA is evident across a wide range of settings of the parameter *α*, while *ρ*PCA is parameter free. This simulated example is typical in biological datasets, where target data includes background noise; in fact, this connection is made explicit in probabilistic linear discriminant analysis (Ioffe, 2006), where the background noise (within cluster in the case of LDA) is an addition to the latent model for the target (between cluster in the case of LDA).

**Fig. 2.**
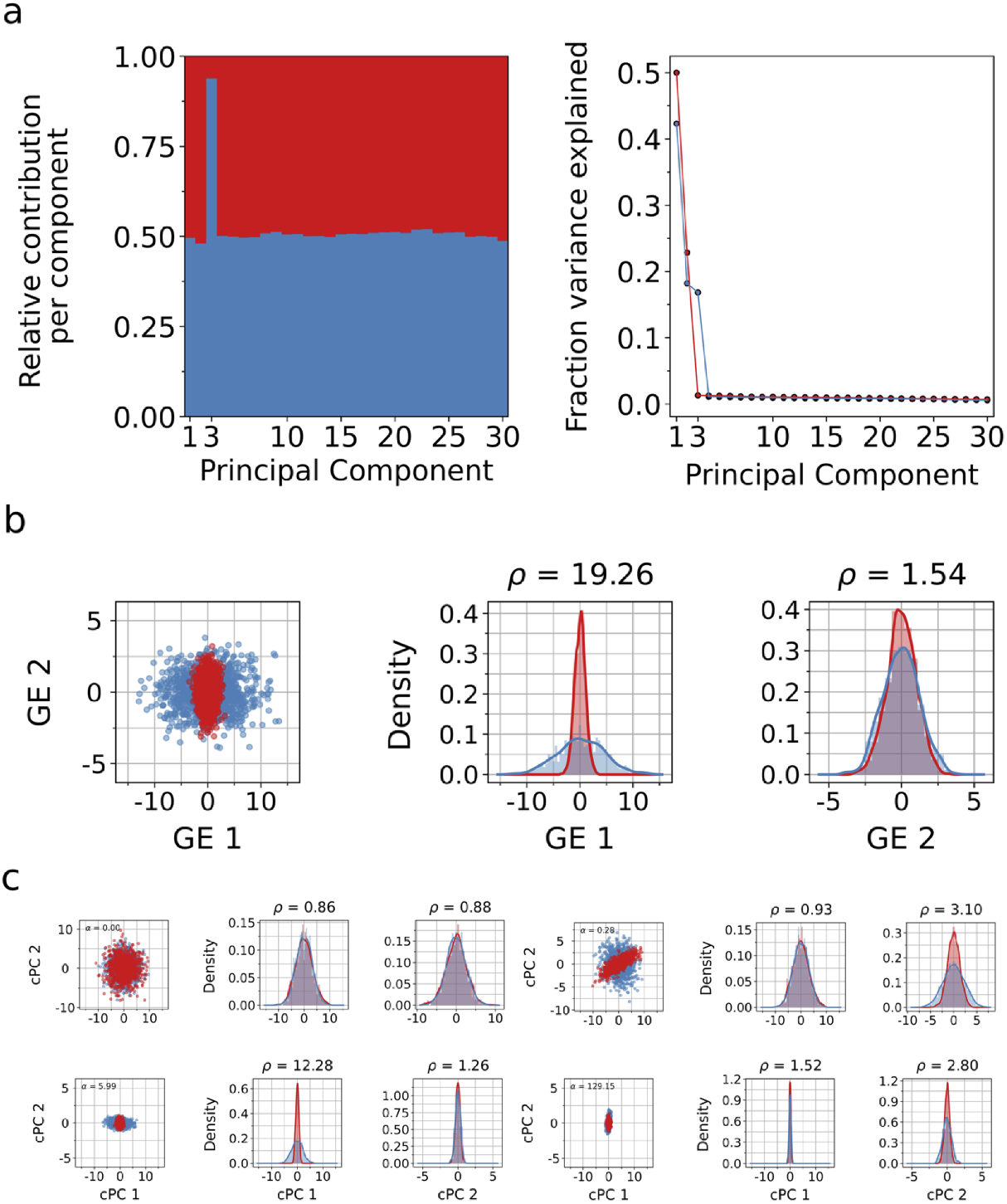
*ρ*PCA correctly identifies orthogonal axes of variation. We simulated 2,000 points in ℝ^30^, where the target and background datasets share variance along two principal directions, while the target exhibits additional variance along a third orthogonal direction (a, left). The remaining dimensions consist of noise with similar variance across both datasets (a, right). *ρ*PCA successfully recovers the single contrastive direction unique to the target, while the remaining generalized eigenvectors capture shared variance between the target and background (b). In contrast, cPCA produces four results corresponding to different values of the hyperparameter *α*. Without prior knowledge of the data structure, it is unclear which choice of *α* yields the correct contrastive component.

Moreover, due to the computational challenge of scanning for an effective parameter setting in cPCA, the implementation of cPCA by (Abid et al., 2018) includes heuristics that further degrade performance. When solving for a problem with 1, 200 features, the (Abid et al., 2018) implementation of cPCA first performs PCA to dimension 1, 000, resulting in a poor projection (Supplementary Fig. 1).

To directly compare cPCA to *ρ*PCA on biological data, we applied *ρ*PCA to a dataset consisting of protein expression measurements from mice exposed to shock therapy (Abid et al., 2018), some of which developed Down Syndrome (DS) and some of which did not (non-DS). As reported in (Abid et al., 2018), performing PCA on the shock-treated mice data does not reveal distinct protein expression between those that developed Down Syndrome and those that did not. Introducing as a background protein expression measurements from non-DS mice that had not received shock therapy, *ρ*PCA finds an effective projection without any parameter tuning (Fig 3). cPCA projections reported in (Abid et al., 2018) as well as those obtained when running cPCA with default parameters produce different projections at different values of *α*, without clear guidance given on how to choose the best value. Moreover, in an improvement with respect to cPCA, the *ρ*PCA projection shows different protein expression patterns between the DS and non-DS mice (Fig 3).

**Fig. 3.**
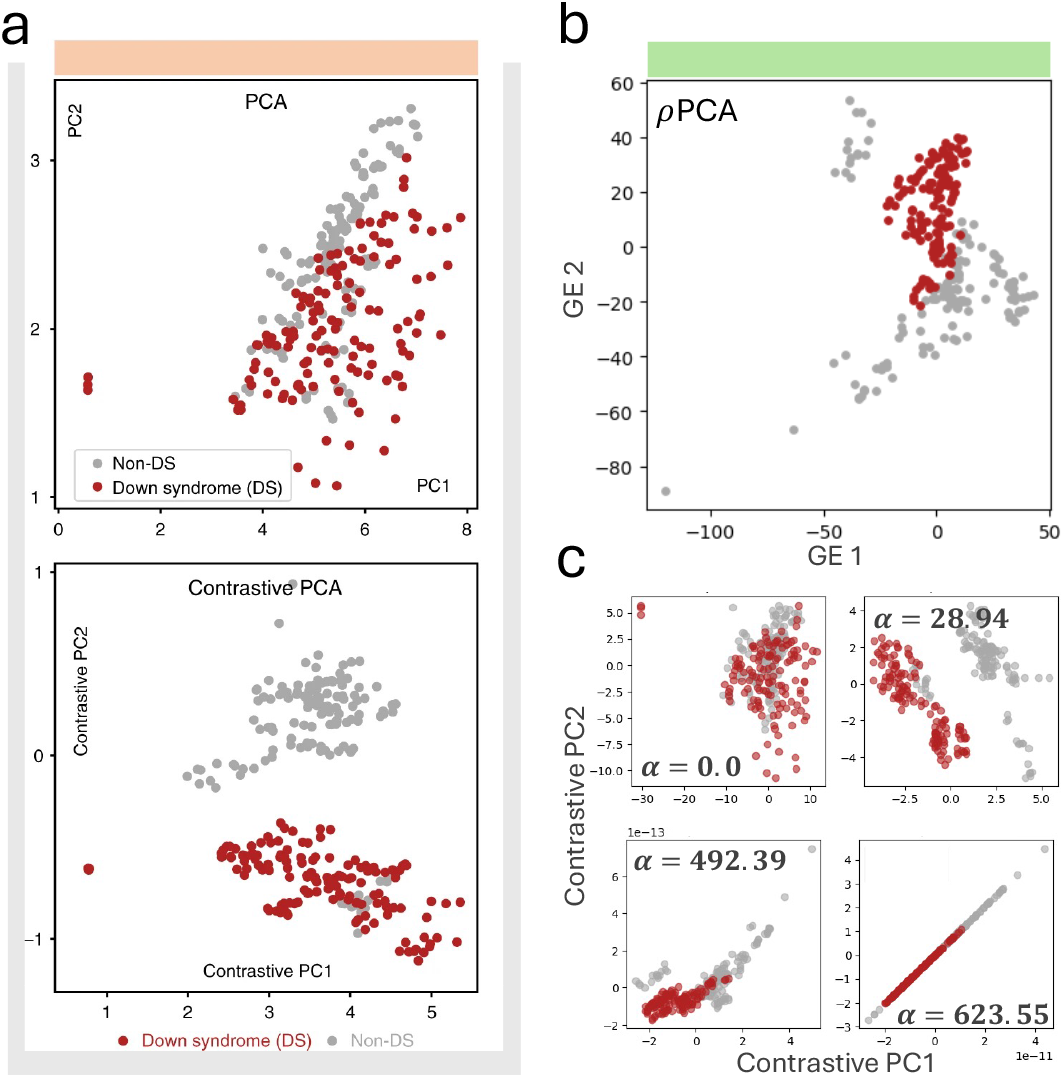
*ρ*PCA recovers structure in biological subgroups without requiring parameter tuning or prohibitively long runtimes. **a**, Original Fig. 3a from (Abid et al., 2018) showing PCA and cPCA projections of protein expression from mice exposed to shock therapy, some of which developed Down Syndrome (DS) and some of which did not (non-DS). **b**, *ρ*PCA recovers separation of the two subgroups without searching over the contrastive parameter *α*. **c**, cPCA by default outputs projections at four values of *α*, with no guidance on how to select the best value.

To demonstrate the interpretability and scalability of *ρ*PCA, we performed an analysis of a large single-nucleus RNA-seq dataset profiling kidney in mice (Rebboah et al., 2025). We applied *ρ*PCA to a target dataset of four female mice with a “background” of four male mice in order to identify cell type specific variation in female gene expression that is distinct from the variation in males. After filtering out low quality nuclei and selecting highly expressed and highly variable genes dataset (see Methods), we analyzed 2,000 genes from 47,798 female nuclei and 42,142 male nuclei of 17 different cell types. While a typical analysis might focus on genes that are differentially expressed (mean or rank differences) between male and female cell types, *ρ*PCA provides a new interpretation of sexually dimorphic gene expression as genes that are *variable* primarily in the target group (female) and not in the background group (male).

The coefficients (loadings) of *ρ*PCA generalized eigenvectors are analogous to PCA loadings, but in this case highlight genes displaying variation among females that is absent from males (Fig. 4). These genes are not identified in a standard application of PCA: unsurprisingly, the genes with the highest loadings on GE 1 and GE 2 are not the genes with the highest loadings on PC 1 and PC 2 (Fig. 4a).

**Fig. 4.**
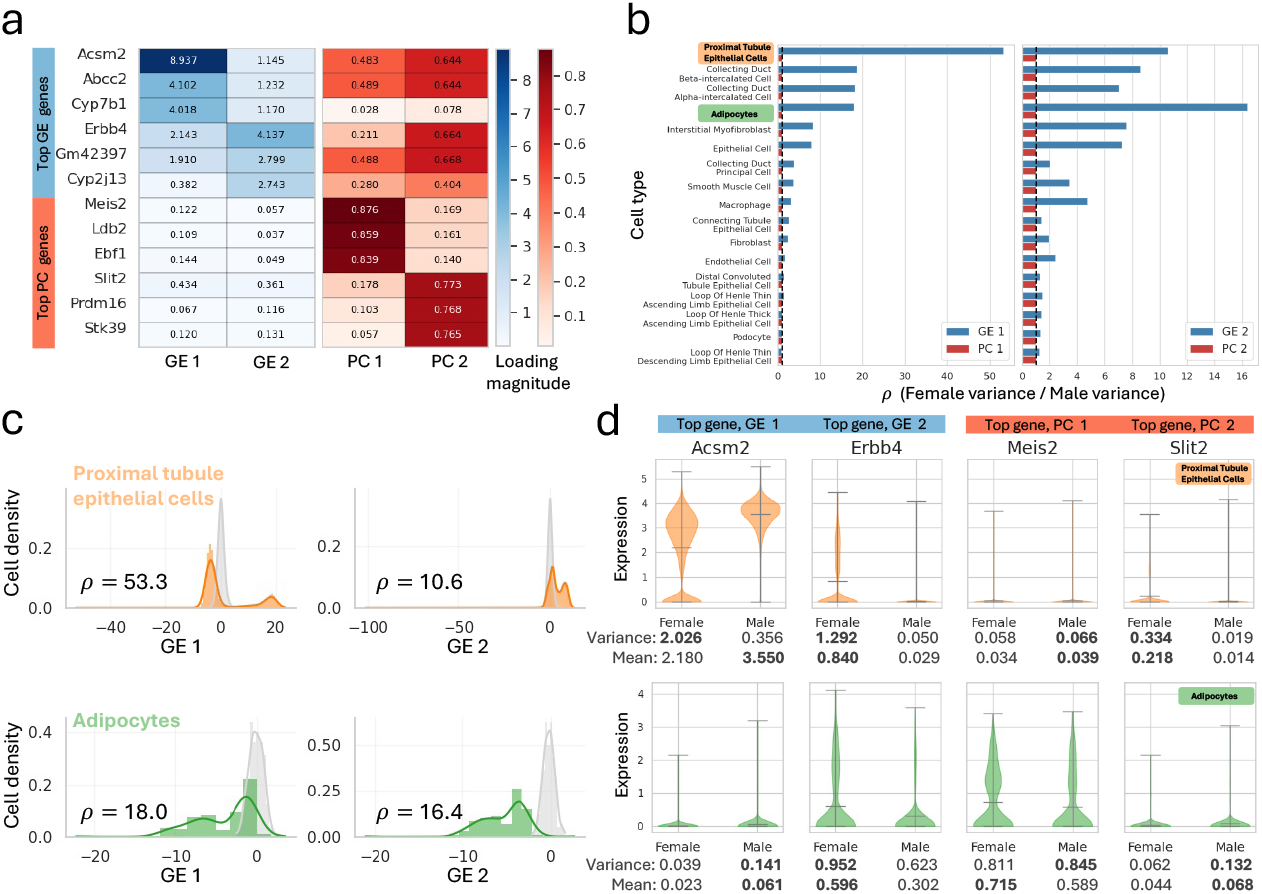
*ρ*PCA analysis of snRNA-seq data from male and female mice reveals genes driving female-specific variation across cell types that PCA fails to identify. **a**, Loadings for the three genes with the greatest magnitude loadings on the first and second generalized eigenvectors (GE 1 and GE 2 respectively) and the first and second principal components (PC 1 and PC 2 respectively): *ρ*PCA highly weights different genes than PCA. **b**, The ratios of variance of female to male nuclei projected onto the first and second GEs for 16 cell types in the kidney. GE 1 separates female proximal tubule epithelial cells, while GE 2 has the highest variance ratio for adipocytes. **c**, Histogram of male and female proximal tubule epithelial cells (orange) and adipocytes (green) projected onto the first and second generalized eigenvectors. **d**, *Acsm2* (gene with the highest magnitude on GE 1), *Erbb4* (gene with the highest magnitude on GE 2), *Meis2* (gene with the highest magnitude on PC 1), and *Slit2* (gene with the highest magnitude on PC 2) expression in male and female nuclei, separated by cell type (proximal tubule epithelial cells, top row, and adipocytes, bottom row).

The variance of projected samples onto each generalized eigenvector also yields insights into the particular cell type driving the main axes of female specific variation. We projected male and female nuclei onto the first and second GEs and calculated the variance per cell type and sex (for all cell types, see Supplementary Fig. 2). We then took the ratio of female to male variance, which is the Rayleigh quotient *ρ* of each GE. We find that GE 1 has the highest female to male Rayleigh quotient in proximal tubule (PT) epithelial cells, while GE 2 has the highest ratio in adipocytes (Fig. 4b,c, with absolute variances shown in Supplementary Fig. 3). As the first generalized eigenvector is the axis that explains the most variation in the target after having accounted for variation in the background, this suggests that of the tested cell types, females show the most unique variability in PT epithelial cells. This is consistent with previous studies: the PT is known to show strong sexually dimorphic gene expression patterns, reported to be regulated by hormone receptors (androgen and estrogen) (Xiong et al., 2023; Ransick et al., 2019).

As expected, the male and female specific expression patterns for the genes driving GE 1 and GE 2 show strong differences in variance. The counts distributions for *Acsm2* (top gene for GE 1) and *Erbb4* (top gene for GE 2) are indeed more variable in females than in males in PT epithelial cells and adipocytes, respectively (Fig. 4d), while the top genes for PC 1 (*Meis2*) and PC 2 (*Slit2*) do not show clear differences in variation. It is interesting that while *Acsm2* is more variable in females than in males in the PT, it has a lower mean expression in females than males. While this gene has been previously noted to have male-biased expression in PT cells (Watanabe et al., 2020), *ρ*PCA identifies that it displays female specificity in the second moment. This variance across cells in females is interesting in light of (Chen et al., 2025), which suggests that *Acsm2* is controlled by androgen in PT cells; each cell may be responding to differences in hormone concentration across the PT. While *ErbB4*, the gene that codes for the receptor protein-tyrosine kinase with roles in cell signaling and growth, is not as well studied, another growth-regulating receptor of the *ErbB* family, *ErbB1*, is known to have sex-biased expression in the kidney and to be sensitive to changes in sex hormones (Zhang et al., 2019). These examples illustrate the potential of *ρ*PCA as a computational approach for identifying genes whose cell-to-cell variation may have functional significance (Dueck et al., 2016).

## Discussion

Subsequent to the introduction of cPCA in (Abid et al., 2018), several variations and derivatives have been proposed (Hawke et al., 2025). For example, (Fujiwara et al., 2019) develop visualization approaches based on cPCA, (Boileau et al., 2020) introduce a sparse version of cPCA, (Fujiwara and Liu, 2023) modify cPCA for binary, ordinal, or nominal data in a method they call cMCA, (Li et al., 2024) develop a probabilistic model whose maximum-likelihood solution yields cPCA in the small variance limit, and (Zhang and Li, 2025) propose a contrastive functional PCA that is cPCA with a weighted covariance matrix. (de Oliveira et al., 2025) introduce six variations of contrastive PCA which are called gcPCA v2.0, v2.1, v3.0, v3.1, v4.0, and v4.1. The gcPCA v2.0 and v3.0 variations are equivalent to each other, the only difference being a shift in eigenvalues. Both are equivalent to the unregularized cPCA^*\**^ of (Golkar et al., 2023) and to *ρ*PCA, although some important mathematical, statistical, and computational motivation for *ρ*PCA are missing from (de Oliveira et al., 2025). As we have shown, the insight that cPCA is an approximation (with heuristics) of *ρ*PCA means that gcPCA v2.0 is not better than cPCA (which (de Oliveira et al., 2025) call gcPCA v1.0) merely because it removes the parameter *α* as suggested in (de Oliveira et al., 2025). Rather, *ρ*PCA is useful in cases where cPCA fails with respect to all parameters *α* (Fig. 2), and even when cPCA can produce a reasonable result, *ρ*PCA is more accurate (Fig. 3). This means that the derivatives of cPCA mentioned above are sub-optimal, in the sense that the cPCA method on which they rely is a heuristic approximation of the Rayleigh quotient solution. Moreover, unlike *ρ*PCA, the gcPCA v4.0 method of (de Oliveira et al., 2025) does not satisfy monotonicity. The gcPCA v2.1, v3.1 and v4.1 variations involve adding an orthogonality constraint as done in (Wang et al., 2018), which as pointed out in (Woller et al., 2025) is problematic.

Indeed, it is no accident that the *ρ*PCA objective has been used for dimension reduction since the introduction of LDA in 1936 (Fisher, 1936), and has been rediscovered in various fields. For example, the extraction of common spatial patterns method (CSP) of (Koles et al., 1990) at its core involves maximizing a Rayleigh quotient, where the target and background covariance matrices are derived from two distinct signal conditions (Cohen, 2022). The resulting generalized eigenvectors define spatial filters that maximize variance in one condition while minimizing variance in the other. CSP has become one of the most popular algorithms for brain computer interface design (Lotte and Guan, 2010). One reason that maximizing the *ρ*PCA objective works well in practice is that *ρ*PCA can be viewed as a restricted Gaussian maximum-likelihood estimator analogous to classical LDA (Hastie and Tibshirani, 1996; Ioffe, 2006). In this view, target and background data arise from Gaussians whose covariances differ by a structured latent component, and the *ρ*PCA solution is the large-sample maximum-likelihood solution. This probabilistic interpretation differs fundamentally from the contrastive latent variable model of Severson et al. (2019), in which both target and background distributions are generated by shared and condition-specific latent variables and the full joint likelihood is maximized. The models of (Hastie and Tibshirani, 1996) and (Ioffe, 2006) provide asymptotic insight into *ρ*PCA, in contrast to the finite-sample likelihood maximization of probabilistic PCA (Tipping and Bishop, 1999). Moreover, the expectation–maximization (EM) algorithm of (Ioffe, 2006) offers an analog of the EM algorithm of (Roweis, 1997) for probabilistic PCA.

The *ρ*PCA method is also powerful due to its efficiency and scalability, making it much more suited to genomics applications than cPCA. We found that *ρ*PCA is much faster than cPCA. For example, in an analysis of a dataset with 10,000 samples and 1,000 features, cPCA takes almost two minutes to run, while *ρ*PCA runs in only a few seconds on the same machine, a speedup of two orders of magnitude. The difference in runtime between *ρ*PCA and cPCA is even more dramatic as the number of samples and features increases (Fig. 5). Analyzing data at increasingly large scales, such as single-cell RNA sequencing data with hundreds of thousands, or even millions, of cells and genes, would require prohibitively long run times with cPCA.

**Fig. 5.**
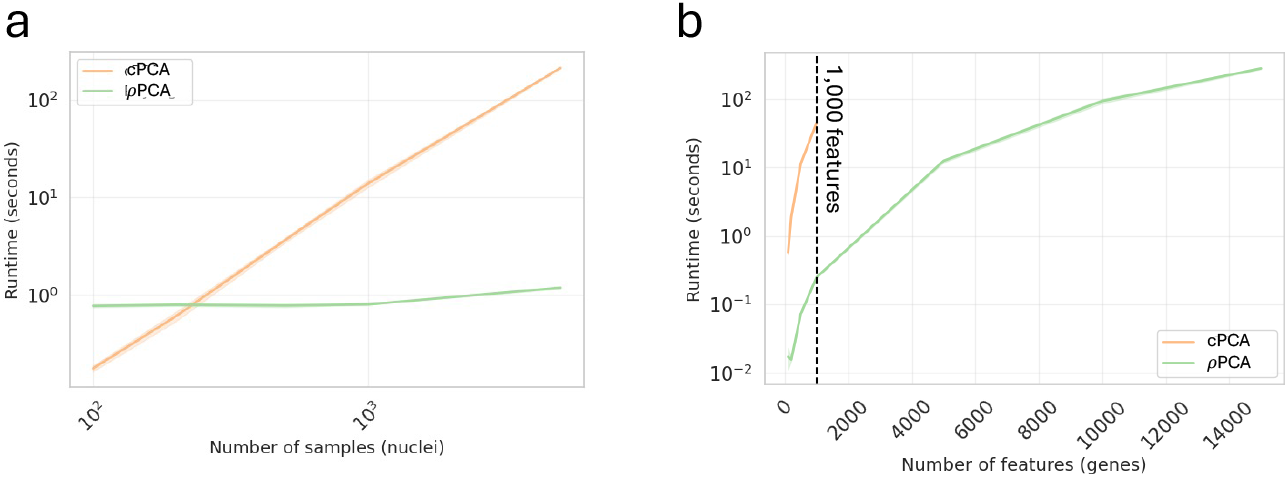
We use snRNA-seq data from mouse kidney to illustrate that *ρ*PCA is orders of magnitude faster than cPCA and remains efficient as **a**, number of samples (with fixed 2,000 features) and **b**, number of features (with fixed 1,000 samples in target and 1,000 samples in background) grow. Note that when samples have more than 1,000 features, cPCA first performs PCA to restrict the number of features to 1,000 (black dashed line).

In summary, the Rayleigh quotient provides an effective unifying principle for linear contrastive learning. This observation can lead to a consolidation of contrastive methods across fields.

## Acknowledgments

Thanks to members of the Pachter lab who provided valuable input in discussions related to the methods during the journal club that led to the work on this paper. MC was funded, in part, by the NSF GRFP under grant no. 2139433. KJ and LP were funded, in part, by NIH 1R01DK143671-02.

## Author contributions

This work emerged as a result of discussions during a journal club on the (Abid et al., 2018) paper organized by MC and KJ. MC, KJ, and LP developed the methods, produced the results and drafted the manuscript.

## Methods

### Nomenclature

The Rayleigh quotient (2) is frequently referred to as *ρ*. We therefore adopt the term *ρ*PCA to highlight the fact that *ρ*PCA extends PCA via a Rayleigh quotient where the matrix *B* in (2) is not necessarily the identity. Furthermore, the term *ρ*PCA rather than just *ρ* is intended to convey that the method is more than just the solution of (5) by computing generalized eigenvalues and eigenvectors of covariance matrices derived from data; *ρ*PCA includes the projections of the data onto the subspaces spanned by the generalized eigenvectors.

Throughout, sample covariance matrices are denoted by 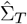 and 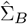. In the large-sample limit *n*_*T*_, *n*_*B*_ → ∞, 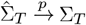 and 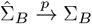, where Σ_*T*_ and Σ_*B*_ are the population covariances.

### Properties of the Rayleigh quotient

If 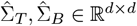 are symmetric matrices with 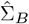 positive definite, *ℛ*_*ρ*PCA_(*v*) satisfies several useful properties. In particular, it is scale invariant: scaling either covariance matrix by a positive constant rescales all eigenvalues but leaves the eigenvectors, and therefore the contrastive subspace, unchanged, i.e., for *a, b >* 0,

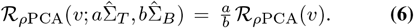

Furthermore, *ρ*PCA satisfies monotonicity: if 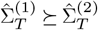 or 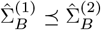, then 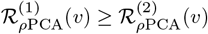. The ⪰ ordering denotes the Loewner (semidefinite) order, where *A ⪰ B* when *A − B* is positive semidefinite. This means that increasing the target variance or decreasing the background variance never decreases the Rayleigh quotient. The method is also rotationally equivariant, in the sense that if both datasets are rotated by the same orthogonal transformation *R*, then the generalized eigenproblem 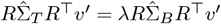 yields transformed solutions *v*^*′*^ = *Rv*, so that the resulting subspace is geometrically consistent under rotation. Finally, the eigen-vectors obtained from *ρ*PCA are mutually orthogonal with respect to the background covariance, satisfying 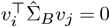 for *i ≠j*, although they may not be orthogonal in the Euclidean inner product. For further discussion of the algebraic properties of generalized Rayleigh quotients, see (Horn and Johnson, 2012) .

### Replicates

In the case of independent, centered, replicate target matrices with the same number of samples *T*_1_, …, *T*_*r*_ and/or background matrices *B*_1_, …, *B*_*s*_ with the same number of samples, *ρ*PCA can be applied to the mean covariance as follows:

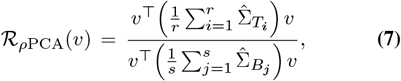

where 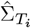 and 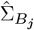 denote the sample covariance matrices of each replicate target and background dataset, respectively.

Averaging the covariance matrices across replicates (or taking the weighted average based on sample size or confidence) is natural because each 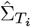 and 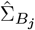 represents an independent empirical estimate of the same underlying covariance covariance matrix (for the target and background conditions, respectively). This therefore increases the effective sample size and reduces estimation variance. The resulting Rayleigh quotient Eq. (7) compares the expected variance of a projection *v* under the target distribution to that under the background distribution.

Note that differential robust PCA (drPCA; Lengerich and Xing (2019)), a method that is not to be confused with drPCA (Chu, 2022), drPCA (Li et al., 2023) or drPCA (Zheng et al., 2024), none of which are contrastive, instead applies (regularized) PCA to the difference vectors of the target and background datasets. The relevance to replicates is that this is equivalent (in the case where the cross-covariance is negligible) to PCA on the sum of target and background covariance matrices. That is, drPCA is PCA applied to 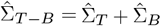 . This same problem afflicts differential PCA (Ji et al., 2013; Yen and Kellis, 2015). In other words, by treating the target and background as if their samples were replicates drawn from the same distribution, differential (robust) PCA becomes the opposite of differential.

### Solving *ρ*PCA

The *ρ*PCA objective (5) seeks to find

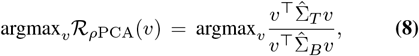

where 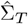 and 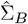 denote the empirical target and background covariances, respectively, and 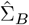 is assumed to be positive definite. That is, a direction *v* in which the target variance is maximized while the background variance is minimized. Because the quotient is homogeneous in *v*, scaling *v* by any nonzero constant does not change the objective. Therefore, we can impose a normalization constraint, leading to the equivalent constrained optimization problem

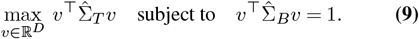

The constraint specifies unit variance in the background, so that the maximization measures relative rather than absolute variance. To solve this constrained problem, we introduce a Lagrange multiplier *λ* and define the Lagrangian

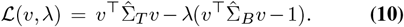

Taking the gradient with respect to *v* and setting it equal to zero yields

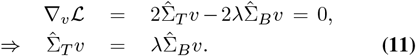

Equation (11) is a generalized eigenvalue problem. Its solutions (*v, λ*) satisfy the stationarity condition of the Rayleigh quotient: the eigenvectors *v* corresponding to the largest generalized eigenvalues *λ* define the directions along which the target variance is maximized relative to the background variance. Each eigenvalue *λ*_*i*_ represents the ratio of target-to-background variance along its associated direction *v*_*i*_.

This extends naturally to multiple dimensions. Let *V* = [*v*_1_, …, *v*_*d*_] ∈ℝ^*D×d*^ denote the matrix whose columns are the top *d* generalized eigenvectors of 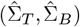, and let Λ = diag(*λ*_1_, …, *λ*_*d*_) be the diagonal matrix of the associated eigenvalues. The multidimensional generalization of the Rayleigh quotient replaces the scalar ratio by a trace ratio, yielding the objective

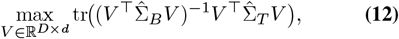

which is equivalent to maximizing 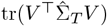 subject to the constraint of background orthonormality 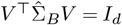. Introducing a symmetric matrix of Lagrange multipliers Λ, the stationarity condition of the Lagrangian

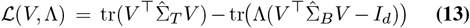

with respect to *V* gives

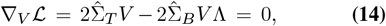

or equivalently,

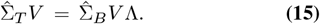

This is the matrix form of the generalized eigenvalue equation: each column of *V* is an eigenvector of 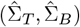 and each diagonal entry of Λ its corresponding eigenvalue. The optimal *V* consists of the *d* generalized eigenvectors associated with the largest eigenvalues, which together span the *ρ*PCA subspace. This subspace maximizes the total target variance relative to background variance, and the constraint 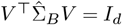 ensures that the learned directions remain mutually orthogonal under the background inner product, 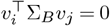 for *i ≠j*.

The reduction of (8) to a generalized eigenvalue problem via the variational argument (9,10,11) as outlined above is the classic Courant-Fischer theorem (Fischer, 1905; Courant, 1920), albeit the Courant-Fischer theorem was initially formulated for PCA (for a history of eigenvalue computation see (Golub and Van der Vorst, 2000)). We have included it for completeness; a recent comment on (de Oliveira et al., 2025) notes that gcPCA v2.0 is equivalent to a generalized eigenvalue problem (Woller et al., 2025), although without reference to the Courant-Fischer theorem.

### Regularization

For simplicity, when describing *ρ*PCA we have assumed that the background covariance matrix 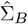 is positive definite, i.e. 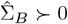. In practice, 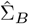 may only be positive semidefinite, i.e., it may not be invertible due to some eigenvalues being equal to zero. This will happen, for example, when there are more features than points in the background data matrix *B*. In such a case, regularization can be used to obtain a well-posed generalized eigenvalue problem. The standard approach to regularization is additive Tikhonov shrinkage, known as ridge regression in the case of least squares problems but used in a similar way for Rayleigh quotient problems (Ye et al., 2006). In additive Tikhonov regularization, the matrix *B* is replaced with 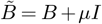 where *µ* ≥ 0 is a parameter. This ensures that the background keeps its original scale, and the regularization is spectrally smooth, i.e. small eigenvalues receive relatively large increases but large eigenvalues are perturbed only a little bit. These regularization features are missing in the cPCA^*\**^ regularization method of (Golkar et al., 2023) presented as an improvement of cPCA, in which *B* is modified as (1 − *β*)*I* + *βB* where 0 ≤ *β* ≤ 1, a method that will change the total variance scale, lead to uniform compression of eigenvalues rather than just lifting small ones, and also loses the standard interpretation of Tikhonov regularization as a maximum-a-posteriori prior (Engl et al., 1996; Hansen, 1998).

### cPCA as an approximation

Contrastive PCA can be understood as a heuristic approximation to (5). To see this, consider the Rayleigh quotient (5) and let 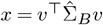. Expanding 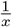 about 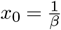 with *β >* 0 gives

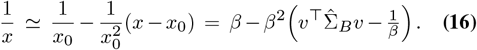

Substituting into *ℛ*_*ρ*PCA_(*v*), disregarding constants, and dividing by 2*β* (which does not change the maximizing direction), yields

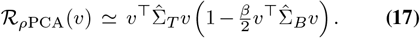

If 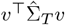 varies slowly in the neighborhood of the maximizing direction, it can be treated as approximately constant, resulting in the cPCA objective

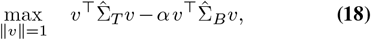

where *α* is a scalar proportional to 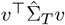, encoding the strength of the background suppression. Thus, the cPCA objective (18) is a first-order heuristic approximation to the *ρ*PCA Rayleigh quotient objective.

### Simulations

To illustrate a failure mode of cPCA, we simulated target and background data from two multivariate Gaussian distributions with mean 0 in 30 dimensions. The target and background data varied along two shared principal directions, while the target had an additional orthogonal axis of variation that was absent in the background. We sampled 1,000 points for the target and another 1,000 points for the background, and then ran *ρ*PCA on the target and background empirical covariance matrices using `scipy.linalg.eigh` (SciPy v1.16.2; Virtanen et al. (2020)) to get generalized eigenvectors and to project the data onto the first and second generalized eigenvectors. We then ran cPCA in default mode (contrastive package v1.2.0) `fit_transform`, which returned results for *α* values of 0.0, 0.28, 5.99, and 129.15.

### Protein expression measurements from shock-treated mice

To show the utility of *ρ*PCA on a dataset for which cPCA was tested in the (Abid et al., 2018) paper, measurements of protein expression in mice subjected to shock therapy (Ahmed et al., 2015; Higuera et al., 2015) were downloaded from the GitHub repository associated to (Abid et al., 2018). We then ran the default `fit_transform` function of cPCA’s contrastive package (v1.2.0) with the shock-treated mice as target samples and the control mice as background samples, which returned projections and visualizations at four alphas (0.0, 28.94, 492.39, and 623.55). We standard scaled the target and background mice matrices separately to column mean of 0 and column variance of 1 before calculating the covariance matrices: after standard scaling, covariance matrices are correlation matrices and performing *rho*PCA can be interpreted as finding sets or modules of correlated genes. We then implemented *ρ*PCA (solving 15) using `scipy.linalg.eigh` (SciPy v1.16.2; Virtanen et al. (2020)).

### snRNA-seq data from mouse kidneys

We obtained processed snRNA-seq counts from the kidney with cell type annotations for 64 mice from (Rebboah et al., 2025). We restricted our analysis to 8 C57BL/6J mice (4 males and 4 females) and the 17 cell types that were not annotated as low quality, resulting in 47,798 nuclei from female mice and 42,142 nuclei from male mice. We next filtered for high expression genes by taking those above the 80th percentile in total counts (sum across all C57BL/6J male and female nuclei). Nuclei were then depth normalized to 10,000 UMIs and transformed with log1p using Scanpy v1.11.4 (Wolf et al., 2018). Genes were further filtered to the 2,000 most highly variable genes selected using female nuclei (`scanpy.pp.highly_variable_genes` with 2,000 top genes and 20 bins).

Next, we performed PCA and *ρ*PCA. Features (genes) across all female and male nuclei were standard scaled (mean of 0 and unit variance) before PCA, which was implemented using `sklearn.decomposition.PCA` (scikitlearn v1.7.2; Pedregosa et al. (2011)). Before *ρ*PCA, female and male count matrices were standard scaled separately, as their covariance matrices are calculated separately. Again, after standard scaling the covariance matrix is a correlation matrix, and genes that are highly weighted along a given generalized eigenvector are those that are correlated. We then calculated the covariance matrices for female and male nuclei counts and solved the generalized eigenvalue problem (15) with females as the target and males as the background using `scipy.linalg.eigh` (SciPy v1.16.2; Virtanen et al. (2020)).

### Timing

We used the previously processed snRNA-seq dataset (see above) to benchmark timing. We increased the number of nuclei (samples) from 100 to 5,000 with a fixed 2,000 gene (features) and the number of features from 100 to 15,000 with a fixed 1,000 samples in the target and 1,000 samples in the background. As cPCA performs PCA to 1,000 features before running their algorithm if samples are of higher dimension, we only ran cPCA to 1,000 features. Benchmarking was performed on a server with 2x Intel Xeon E5-2690 CPUs (16 cores, 32 threads total) and 128 GB RAM.

## Code and data availability

A pip installable package for performing *ρ*PCA is available, with source and an example use case, at https://github.com/pachterlab/rhopca/. While running *ρ*PCA can be accomplished with one line in Python, the rhopca package facilitates easy use of the method for single-cell genomics by directly enabling analysis of Scanpy anndata objects. All code necessary to generate the schematic (Fig. 1), simulations (Fig. 2) and biological analyses (Figs. 3, 4, and 5) is available at https://github.com/pachterlab/CJP_2025/. Data for Fig. 3 is at https://github.com/pachterlab/CJP_2025/, and data for Fig. 4 and Fig. 1 has been uploaded to Zenodo (DOI: 10.5281/zenodo.17487639).

