## Supplementary Figures for "The Rayleigh Quotient and Contrastive Principal Component Analysis I"

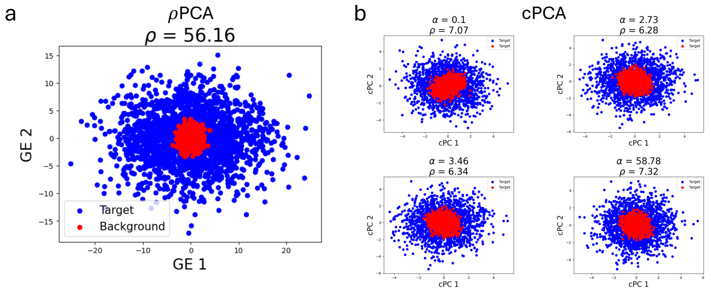


Figure S1: Simulations show that running default contrastive PCA [Abid et al., 2018] does not accurately shrink background variance in part because of the PCA performed when the number of features exceeds 1,000. cPCA first performs PCA to reduce dimensionality of the dataset when the number of features is greater than 1,000. **a,** $\rho$PCA projections of target and background samples produces a $\rho$ (target variance / background variance) of 56.16 along the first generalized eigenvector (GE 1), while **b,** cPCA's largest variance ratio along the first contrastive PC (cPC 1) is 7.32 for the returned projections at default $\alpha$ values. cPCA run with default settings returned projections at $\alpha = 2.73, 3.46$ and 58.78. We also include $\alpha = 0.1$ to show that an arbitrary, smaller value of $\alpha$ has a similar target / background variance ratio to the larger, returned values.

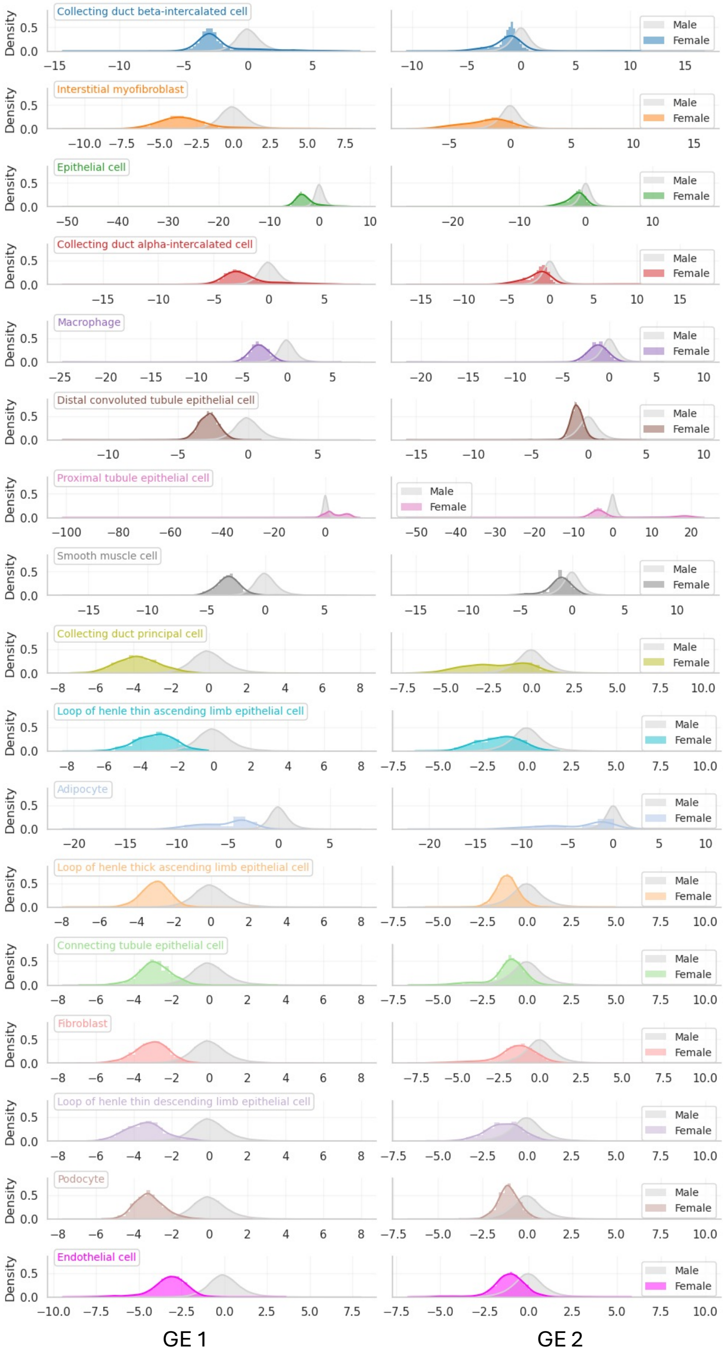


Figure S2: Histograms of projected target (female) and background (male) nuclei onto GE 1 and GE 2, separated by cell type as annotated in [Rebboah et al., 2025]. Data processing and $\rho$PCA fitting described in Methods under subsection "snRNA-seq data from mouse kidneys."

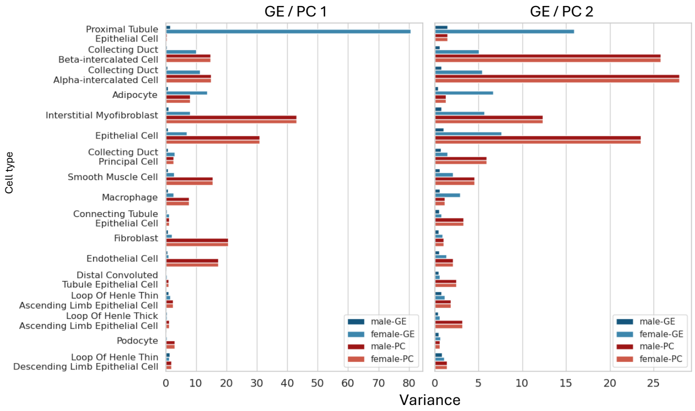


Figure S3: Variance of projected target (female) and background (male) nuclei onto GE 1, GE 2, PC 1 and PC 2, separated by cell type as annotated in [Rebboah et al., 2025]. Data processing and $\rho$PCA fitting described in Methods under subsection "snRNA-seq data from mouse kidneys."

### References

Abubakar Abid, Vivek Zhang, Vibhor Bagaria, and James Zou. Contrastive principal component analysis. *Advances in Neural Information Processing Systems*, 31, 2018.

Elisabeth Rebboah, Ryan Weber, Elnaz Abdollahzadeh, Nikhila Swarna, Delaney K Sullivan, Diane Trout, Fairlie Reese, Heidi Yahan Liang, Ghassan Filimban, Parvin Mahdipoor, et al. Systematic cell-type resolved transcriptomes of 8 tissues in 8 lab and wild-derived mouse strains captures global and local expression variation. *bioRxiv*, pages 2025–04, 2025.
